# Conformation-specific synthetic antibodies discriminate multiple functional states of the ion channel CorA

**DOI:** 10.1101/2023.05.07.539746

**Authors:** Satchal K. Erramilli, Pawel K. Dominik, Dawid Deneka, Piotr Tokarz, Sangwoo S. Kim, Bharat G. Reddy, Blazej M. Skrobek, Olivier Dalmas, Eduardo Perozo, Anthony A. Kossiakoff

## Abstract

CorA, the primary magnesium ion channel in prokaryotes and archaea, is a prototypical homopentameric ion channel that undergoes ion-dependent conformational transitions. CorA adopts five-fold symmetric non-conductive states in the presence of high concentrations of Mg^2+^, and highly asymmetric flexible states in its complete absence. However, the latter were of insufficient resolution to be thoroughly characterized. In order to gain additional insights into the relationship between asymmetry and channel activation, we exploited phage display selection strategies to generate conformation-specific synthetic antibodies (sABs) against CorA in the absence of Mg^2+^. Two sABs from these selections, C12 and C18, showed different degrees of Mg^2+^-sensitivity. Through structural, biochemical, and biophysical characterization, we found the sABs are both conformation-specific but probe different features of the channel under open-like conditions. C18 is highly specific to the Mg^2+^-depleted state of CorA and through negative-stain electron microscopy (ns-EM), we show sAB binding reflects the asymmetric arrangement of CorA protomers in Mg^2+^-depleted conditions. We used X-ray crystallography to determine a structure at 2.0 Å resolution of sAB C12 bound to the soluble N-terminal regulatory domain of CorA. The structure shows C12 is a competitive inhibitor of regulatory magnesium binding through its interaction with the divalent cation sensing site. We subsequently exploited this relationship to capture and visualize asymmetric CorA states in different [Mg^2+^] using ns-EM. We additionally utilized these sABs to provide insights into the energy landscape that governs the ion-dependent conformational transitions of CorA.

## Highlights

- CorA is the primary transport protein in prokaryotes for Mg^2+^, the most abundant divalent cation.
- Conformation-specific sABs probe different features of CorA in open-like conditions.
- Biophysical assays and ns-EM show sAB C18 is highly specific to the Mg^2+^-free state of CorA.
- X-ray structures shows sAB C12 is a competitive inhibitor of regulatory Mg^2+^ binding to CorA.
- ns-EM shows the sABs capture a range of asymmetric CorA conformations in different [Mg^2+^].

## Introduction

Most protein functions are governed by sets of programed conformational transitions, many of which are exceedingly subtle. Further, many of these transitions are transient in nature and attempts to characterize them in any detail have been thwarted by a number of technical challenges. As a result, this leaves researchers to rely on low resolution methods and modeling to infer the structural connections between dynamics and function. To fill this void, we have developed a technology platform and demonstrated its ability to overcome a number of these hurdles[1–4]. A key element of this platform is a phage display methodology using powerful selection strategies to generate tailored Fab-antibody fragments referred to as “synthetic antibodies” or sABs[5–7]. These sABs can have exquisite sensitivity to conformation; hence, to the functional dynamics of proteins[3,8,9]. They have proven to be extremely useful in applications of X-ray crystallography and cryo-electron microscopy (cryo-EM), facilitating the elucidation of structures of important proteins previously resistant to analysis[10–17]. Notably, sABs also offer value as probes in biological assays[18–20].

Over the last several years, this approach has proven transformative for solving numerous high-profile membrane proteins by cryo-EM, exploiting the sABs as high-performance fiducial markers that increase the mass of the particle and aid in image alignment [14–17,21–23]. To further investigate the potential of this approach in stabilizing functionally important conformational states, we chose the pentameric magnesium channel CorA from *Thermatoga maritima* as a model system. CorA is the main magnesium uptake system in bacteria and archaea and works as a Mg^2+^-gated Mg^2+^-selective ion channel[24,25]. In its closed, non-conductive Mg^2+^-bound form, it crystallizes in the presence of non-physiologically high cation concentrations in multiple compact crystallographic lattices as a symmetric pentamer[26–31]. However, in its “open” form, there appears to be no well-defined single conformational state and consequently this state has been recalcitrant to crystallization apparently because it is too dynamic [32,33].

The crystal structure of the closed form along with spectroscopic data revealed that sets of acidic residues function as divalent cation sensors[29,34–36]. These include the D89 and D253 pair, along with E88, D175, and D179; these sets of residues form what are known as the M1 and M2 sensor sites. This suggests that the presence of coordinated divalent cations at these binding sites triggers a conformational change that is transmitted along the pore forming helices constituting the permeation pathway, leading to the non-conductive, closed state. More recently, low resolution (∼7 Å) structures of the Mg^2+^ free “open” state form of *Tm*CorA were determined by cryo-EM, showing multiple asymmetric conformations among the five N-terminal domains[33]. Determining which conformations were conducive to ion permeability was not feasible at this resolution.

We believed that by utilizing a collection of conformationally specific synthetic antibodies (sABs) that we had previously developed for the Mg^2+^-free CorA molecule, we could effectively discern the functional states responsible for ion conductance in CorA[7]. Some of the sABs that we generated demonstrated exceptional specificity for the Mg^2+^-depleted channel (Mg-dep CorA;, 1mM EDTA). We also identified sABs that bound to the channel at both Mg^2+^-depleted conditions and physiological Mg^2+^ concentrations (1-20 mM MgCl_2_), but not at the high Mg^2+^ concentrations used in the X-ray and cryo-EM structures that exhibit 5-fold symmetry. This implied that the closed conformation of CorA under physiological Mg^2+^ concentrations (Mg-phys;, 1-20 mM MgCl_2_) is also asymmetric to a degree and that strict 5-fold symmetry might be an artifact of the high cation concentrations used in the experimental conditions.

Here we further characterize conformation specific sABs that bind and stabilize two functional conformational states: a Mg^2+^-depleted state (sAB C18) and one that exemplifies physiological conditions (sAB C12). We find that sAB C18 traps the Mg-dep state and is highly sensitive to Mg^2+^ concentration, while sAB C12 binds at physiological concentrations (Mg-phys) and competes with Mg^2+^. We observe that sAB binding in the case of the Mg-dep condition results in a 1 to 1 complex between sAB C18:CorA, while for sAB C12, in either depleted or physiological Mg^2+^ conditions, there is a mixture of stoichiometries between the sAB and CorA: 1 or 2 C12 sAB(s) per pentamer. These variable stoichiometries indicates that there are multiple conformations represented in the sAB:CorA complexes at both the Mg-dep and Mg-phys conditions.

To better understand the nature of the competitive binding behavior of C12 with Mg^2+^, we determined the structure of sAB C12 bound to the N-terminal domain (NTD) of CorA at 2.0 Å resolution. The structure reveals that C12 makes extensive interactions with the NTD along an interface that is inaccessible based on the symmetric model of the CorA pentamer (in its putative closed state). Given that the phage display selection conditions used to generate the C12 sAB are highly unlikely to induce a non-native conformation on the channel, we believe the C12 epitope becomes readily available at physiological concentrations of Mg^2+^. The structure also shows how the mode of binding of one of the complementarity determining regions (CDR H3) mimics the binding of regulatory Mg^2+^ ions in the channel and thus provides insight into the competitive binding mode of the sAB.

## Results

### Selection of conformation-specific sABs against CorA

As was previously reported[7], we selected sABs specific to nanodisc-reconstituted wild-type CorA in magnesium-free conditions by simultaneously counter-selecting against the CorA^D253K^ mutant channel in solution (**Fig. 1A)**. This mutation was shown to be insensitive to magnesium in solution and maintains the five-fold symmetric conformation of the cytoplasmic domains irrespective of [Mg^2+^], thus resembling the wild-type closed structure[36,37]. Hence, competition with the mutant channel counter-selected sABs that were conformation agnostic and consequently eliminated from the final pool. We generated twenty-two unique sABs from this selection campaign, most of which were sensitive to environmental MgCl_2_ to varying extents[7]. We selected from this cohort sABs C18 and C12, which both bound Mg-dep CorA with sub-nanomolar affinities based on ELISA measurements (**Fig. 1B**). Single-point ELISA measurements using constant sAB concentrations showed sAB C18 was highly specific to CorA in the Mg-dep state, with negligible binding in the Mg-phys condition, while sAB C12 showed robust but slightly reduced binding to CorA in the Mg-phys state compared with the Mg-dep state (**Fig. 1C**). To further dissect their respective sensitivities to environmental Mg^2+^, we performed surface plasmon resonance (SPR) to measure binding kinetics of C18 and C12 under these different conditions. These experiments elucidated the relationship between CorA magnesium binding and binding of the sABs to their respective epitopes across Mg^2+^ concentrations that encompass the physiologic concentration range. We further characterized binding stoichiometries using negative-stain electron microscopy (ns-EM) to gain insights into conformational transitions involved in CorA ion conductance.

**Figure 1.**
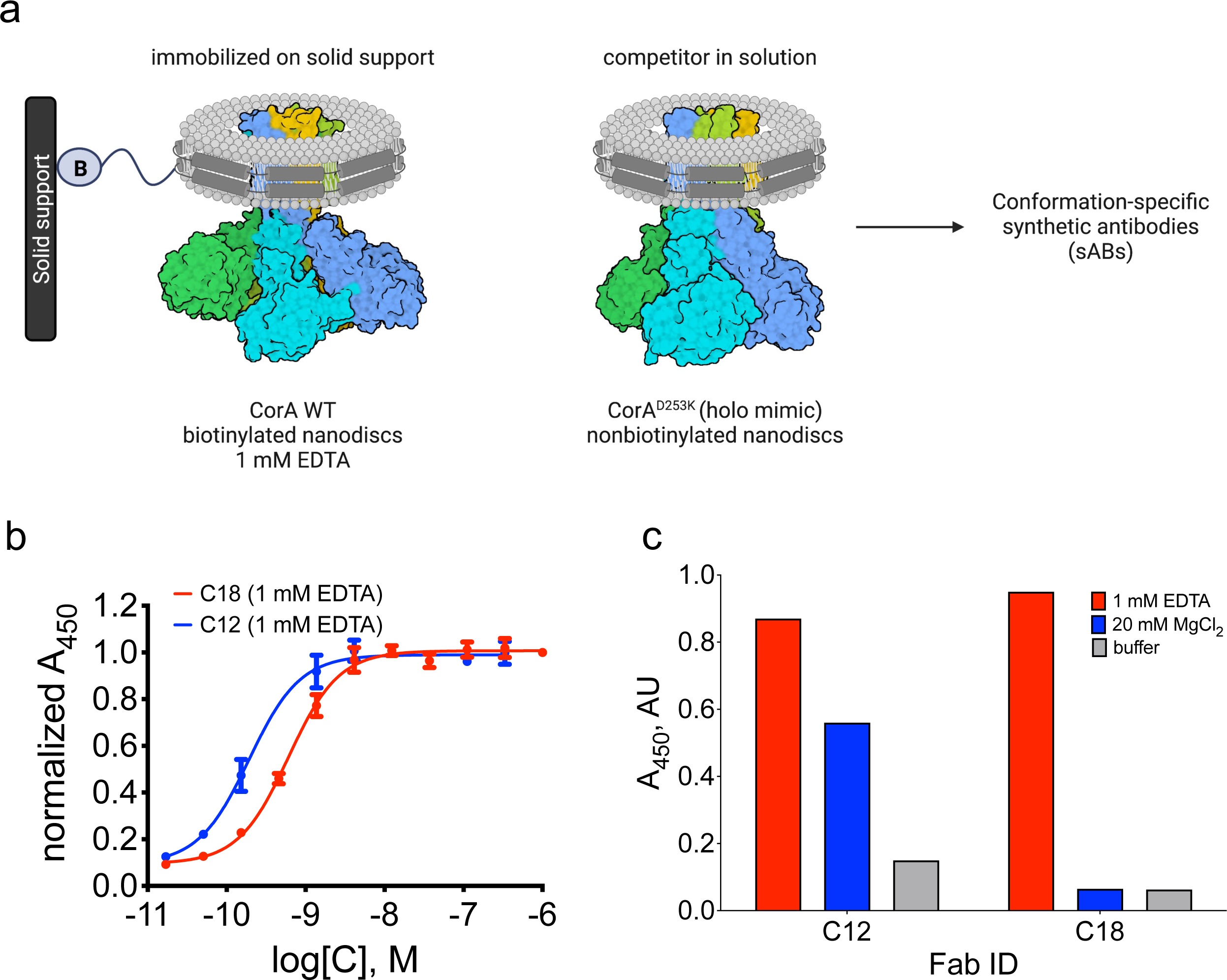
Generation and characterization of conformation-specific sABs against CorA. **A.** Phage display selection strategy for conformation-specific sABs against CorA involved counter selection with CorA D253K, which mimics the magnesium-bound state in the cytoplasmic regulatory domain. **B.** Affinity estimation ELISA of purified Fabs C12 (EC_50_ ∼192 pM) and C18 (EC_50_ ∼610 pM) shows high affinity binding to CorA in 1 mM EDTA. Assays performed in triplicate. Means and error bars plotted together with sigmoidal dose response curve. **C.** Single point binding data of purified Fabs C12 and C18 to CorA in biotinylated nanodiscs immobilized in ELISA wells. Fabs tested in ELISA buffer with 20 mM MgCl_2_ (blue), without (1 mM EDTA, red), or a buffer-only control (gray).

### Conformation-specific sAB C18 binds CorA across a narrow range of Mg^2+^ concentrations bordering on the physiological range

His-tagged DDM-solubilized CorA was immobilized on the SPR chip sensor surface, and binding of the sABs was measured in SPR running buffer with increasing concentrations of MgCl_2_. Kinetic measurements were performed in each of these buffer conditions. The results showed affinities of the Mg-dep specific sAB C18 were dependent on Mg^2+^ concentration within a narrow range. C18 bound CorA with K_D_ values of 11-66 nM in 0-500 µM Mg^2+^ concentrations (**Fig. 2A-C**), with good agreement in affinities with the ELISA data. A sharp transition in K_D_ occurred after this magnesium concentration range, and the affinity was nearly 20 times lower at 750 µM Mg^2+^ (**Fig. 2D**). This precipitous decline in sAB affinity continued with increasing [MgCl_2_], with a ten times lower K_D_ at 1 mM, and no significant binding measured at 1.5 and 2.0 mM Mg^2+^ concentrations (**Fig. 2E-F**). SPR kinetic parameters established this decrease in affinity with increasing Mg^2+^ concentration was primarily due to decreasing on-rates, with relatively small changes to the off-rates in the 0-1.0 mM range (**Fig. 2G, Supplemental Table 1**). The SPR measurements also showed progressive reduction in relative sAB response as the concentration of Mg^2+^ increased. These data demonstrate that C18 can still bind when some magnesium is bound to CorA, but over a short dynamic range near the limits of physiologic Mg^2+^ concentrations. The decrease in on-rates as a function of increasing [Mg^2+^] implies Mg^2+^ has to be displaced for the epitope to be accessible, while the invariant off-rates over the 0-1.0 mM range demonstrate a decreased influence of Mg^2+^ once the sAB is bound. Using these measurements, we determined the free energy difference between the Mg^2+^ bound (1 mM) and depleted (0 mM) CorA states to be 4.1 kcal/mol.

**Figure 2.**
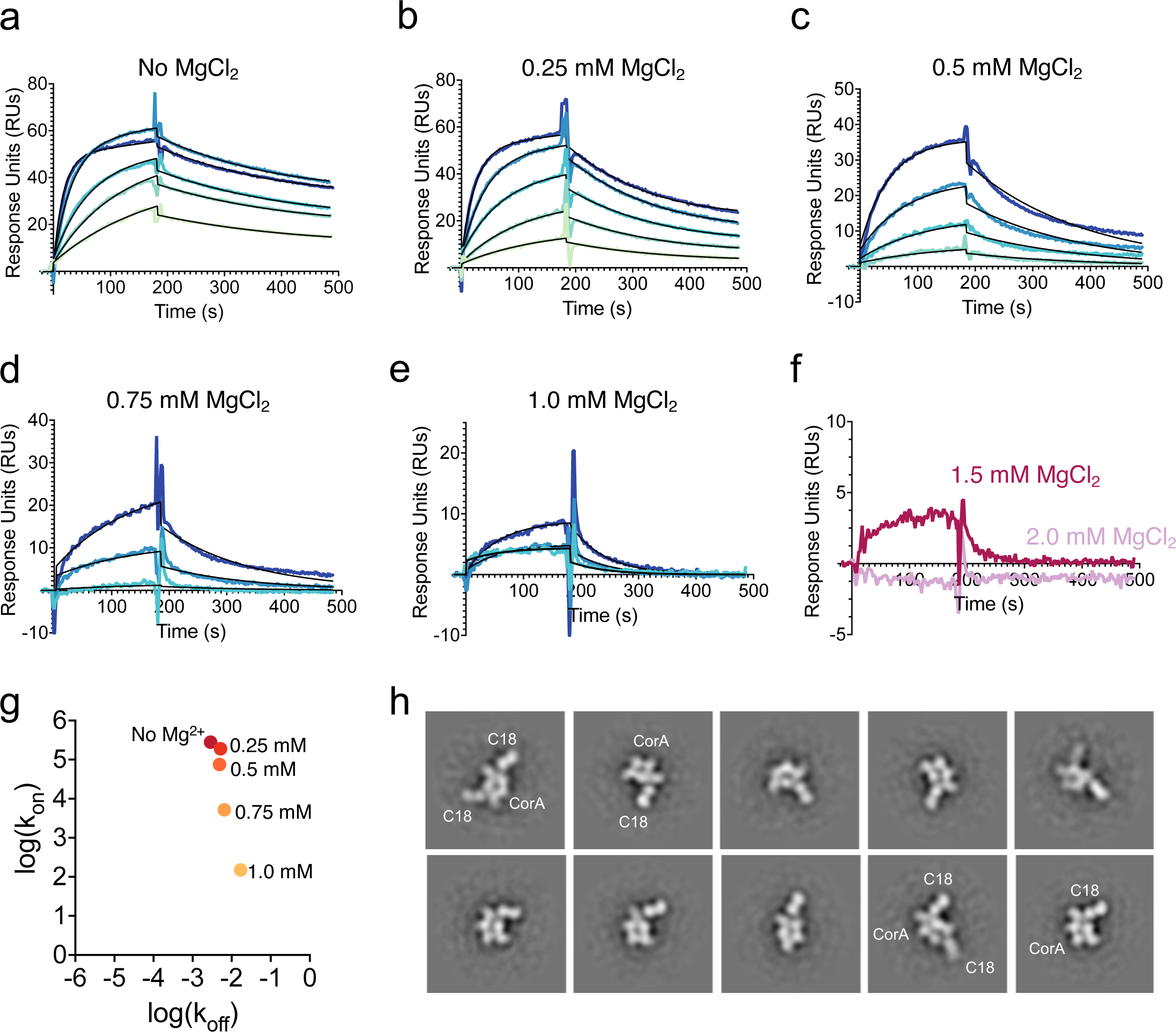
Dissecting the relationship between environmental Mg^2+^ concentration and sAB C18 binding. **A-F**. Double-referenced SPR sensorgrams of a two-fold dilution series of sAB C18 (200-12.5 nM) binding to immobilized CorA in the presence of 0 (A., 11.0 nM K_D_), 0.25 (B., 28.0 nM K_D_), 0.5 (C., 65.8 nM K_D_), 0.75 (D., 1.26 µM K_D_), 1.0 (E., 11.3 µM K_D_), 1.5 (F., dark purple), and 2.0 (F., light purple) mM Mg^2+^. For A-E, experimental curves are overlaid with kinetic fitted curves (black) from a one-to-one binding model. Kinetic fitting was not performed for the last two concentrations (F). **G**. Plot of log(k_on_) vs. log(k_off_) for C18 SPR data from 0-1 mM Mg^2+^. **H**. Negative-stain EM two-dimensional class averages of the nanodisc-reconstituted CorA-C18 complex in 1 mM EDTA. Sample micrograph shown in **Supplemental Figure 1**.

We then used ns-EM to further characterize C18 binding to CorA in 1 mM EDTA. Two-dimensional class averages showed a bimodal population distribution segregated by compositional differences, with either one or two sABs bound per CorA pentamer. The former appears to be the more abundant stoichiometric class. We interpret these results as consistent with the cryo-EM structures and an atomic force microscopy analysis of the Mg-dep condition, which both showed at least two major open conformations at low resolutions[32,33]. This conformational distribution is likely reflected by the compositional differences of the ns-EM data. In combination with the SPR data, these results show the open-like conformations are only present in a narrow range of MgCl_2_ concentrations, with a major structural transition after 0.5 mM making this epitope largely unavailable. C18 thus acts as a probe for regulatory Mg^2+^ binding and the resulting ion-dependent conformational transitions.

### Conformation-specific sAB C12 demonstrates competitive binding behavior with _Mg2+_

We measured C12 binding to immobilized his-tagged CorA by SPR in 0 and 20 mM MgCl_2_, and found a modest change in K_D_, from 0.8 to 5.9 nM (**Fig. 3A, Supplemental Fig. 2A-B, Supplemental Table 1**), with a 1.2 kcal/mol free energy difference between the two conditions. Similar to C18, a decrease in sAB response was observed due to the presence of Mg^2+^, from an R_max_ of 105 to 45 RUs in 0 and 20 mM MgCl_2_, respectively. These measurements suggested a mechanism for environmental Mg^2+^ sensitivity that was different than what was observed for C18.

**Figure 3.**
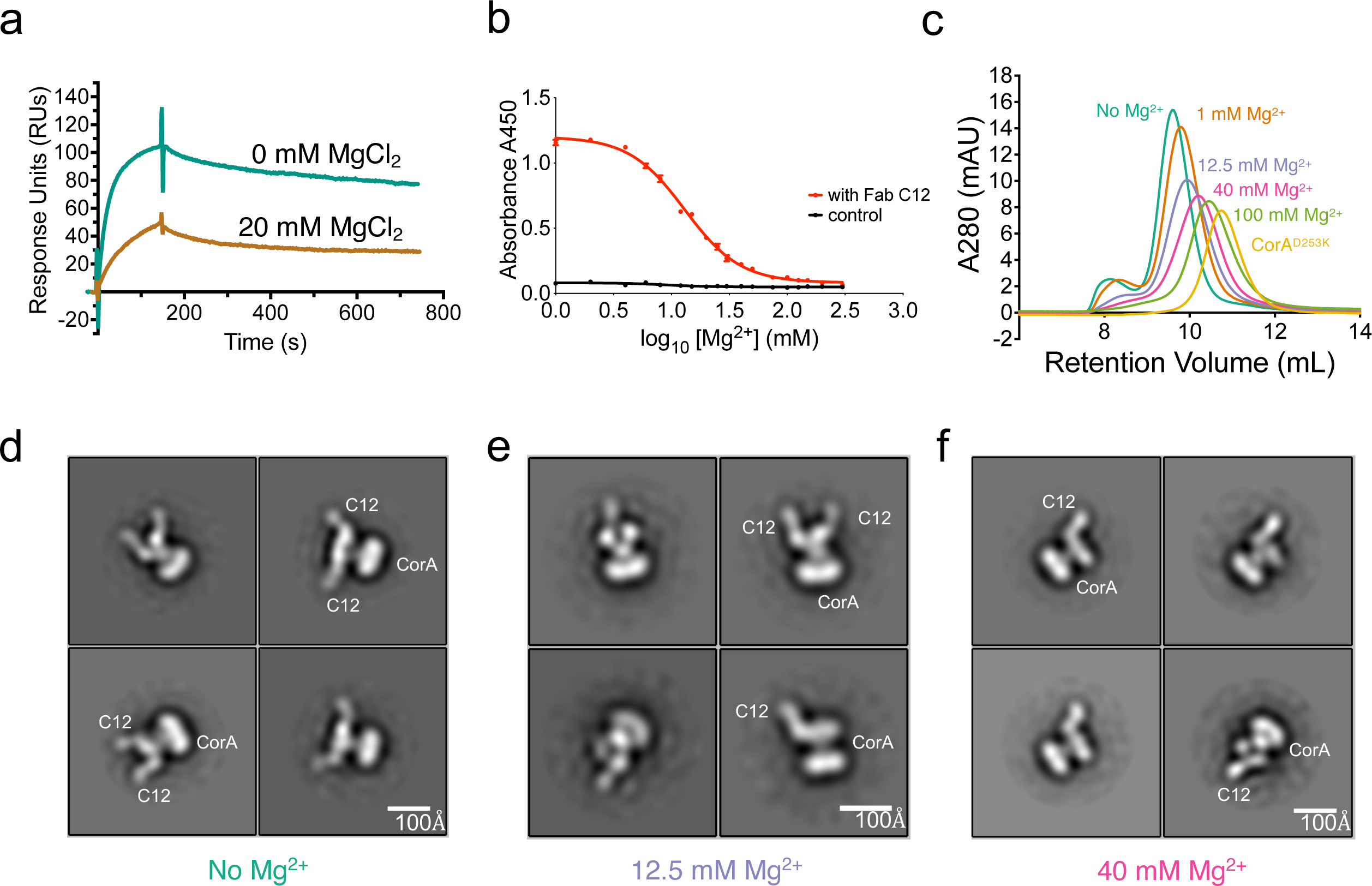
C12 shows a stepwise binding model to CorA in the presence of increasing [Mg^2+^]. **A**. SPR sensorgrams depict 25 nM C12 binding to CorA to determine binding kinetics without Mg^2+^ (teal, 0.83 nM K_D_) and with 20 mM Mg^2+^ (brown, 5.9 nM K_D_). See **Supplemental Table 1** and **Supplemental Fig. 2. B**. ELISA measuring binding of C12 at constant concentration to CorA in different magnesium concentrations. The EC_50_ for Mg^2+^ is 13.03 millimolar, compared to the K_D_ of CorA for Mg^2+^ without sAB C12 (1-2 mM). C. Overlay of size-exclusion chromatograms of CorA-C12 complexes in different environmental magnesium concentrations. Curves are colored and labeled according to condition. D. Negative stain EM 2D classes of CorA-C12 complex in 1 mM EDTA showing two visible sABs. The distance between the two visible sAB lobes in the side view on the bottom right class was ∼150 Å. E. Negative stain EM 2D classes of CorA-C12 in 12.5 mM MgCl_2_. A mixture of classes with one and two bound sABs are visible. The approximate distribution for each was 50/50 from a set of 6500 particles. F. Negative stain EM 2D classes of CorA-C12 in 40 mM MgCl_2_. Only one sAB is visible in views. Sample micrograph in **Supplemental Figure 2**.

To better understand this relationship, we tested C12 binding to CorA by ELISA in the presence of increasing concentrations of Mg^2+^. CorA in biotinylated nanodiscs was immobilized onto ELISA plates in buffer containing increasing concentrations of MgCl_2_ that spanned orders of magnitude below and above the physiological range. A constant concentration of sAB C12 was used to measure binding in each magnesium condition. The results showed concentrations much higher than the typical physiological [Mg^2+^] were required to completely displace sAB C12. We determined an EC_50_ for magnesium binding by CorA of 13 mM (**Fig. 3B**), about ten times higher than the reported K_D_[36]. These data suggested a competitive binding model.

We further characterized C12 using size-exclusion chromatography and negative-stain EM. We tested sAB binding in 0, 1, 12.5, 40, and 100 mM MgCl_2_, as well as to the closed state mimic, CorA^D253K^ (**Fig. 3C**), by adding a two-fold molar excess of C12 per CorA monomer to form fully saturated complexes. We observed several transitions to later retention volumes of CorA-C12 complexes due to the addition of Mg^2+^, and a sharp decrease in A_280_ between 1 and 12.5 mM MgCl_2_. No C12 binding was observed to CorA^D253K^. We then collected ns-EM data for the 1 mM EDTA, 12.5 mM MgCl_2_, and 40 mM MgCl_2_ samples (**Supplemental Fig. 3A-B**). Two-dimensional class averages of these samples showed varying distributions of CorA-C12 complex stoichiometries. The 1 mM EDTA population consisted entirely of CorA with two bound sABs (**Fig. 3D**). The population shifted upon the addition of increasing amounts of Mg^2+^, from a mixture of two-and one-sAB complexes (12.5 mM MgCl_2_, **Fig. 3E**) to entirely one-sAB complexes (40 mM MgCl_2_, **Fig. 3F**). The changes in population stoichiometries were consistent with the shifts in retention volumes observed by SEC. We therefore concluded that the binding stoichiometry was inversely correlated with environmental magnesium concentrations.

### Structure of the complex between the soluble N-terminal domain of CorA and sAB C12

Due to the magnesium sensitivity of C12, we sought to further characterize its binding mode to CorA. We found C12 formed a stable complex with the isolated soluble N-terminal cytosolic sensor domain of CorA (CorA^NTD^; residues 1-280; **Fig 4A**). We were not able to isolate a C18-CorA^NTD^ complex (data not shown), indicating this sAB likely bound to an epitope more evenly distributed across adjacent protomer interfaces in the full-length CorA.

**Figure 4.**
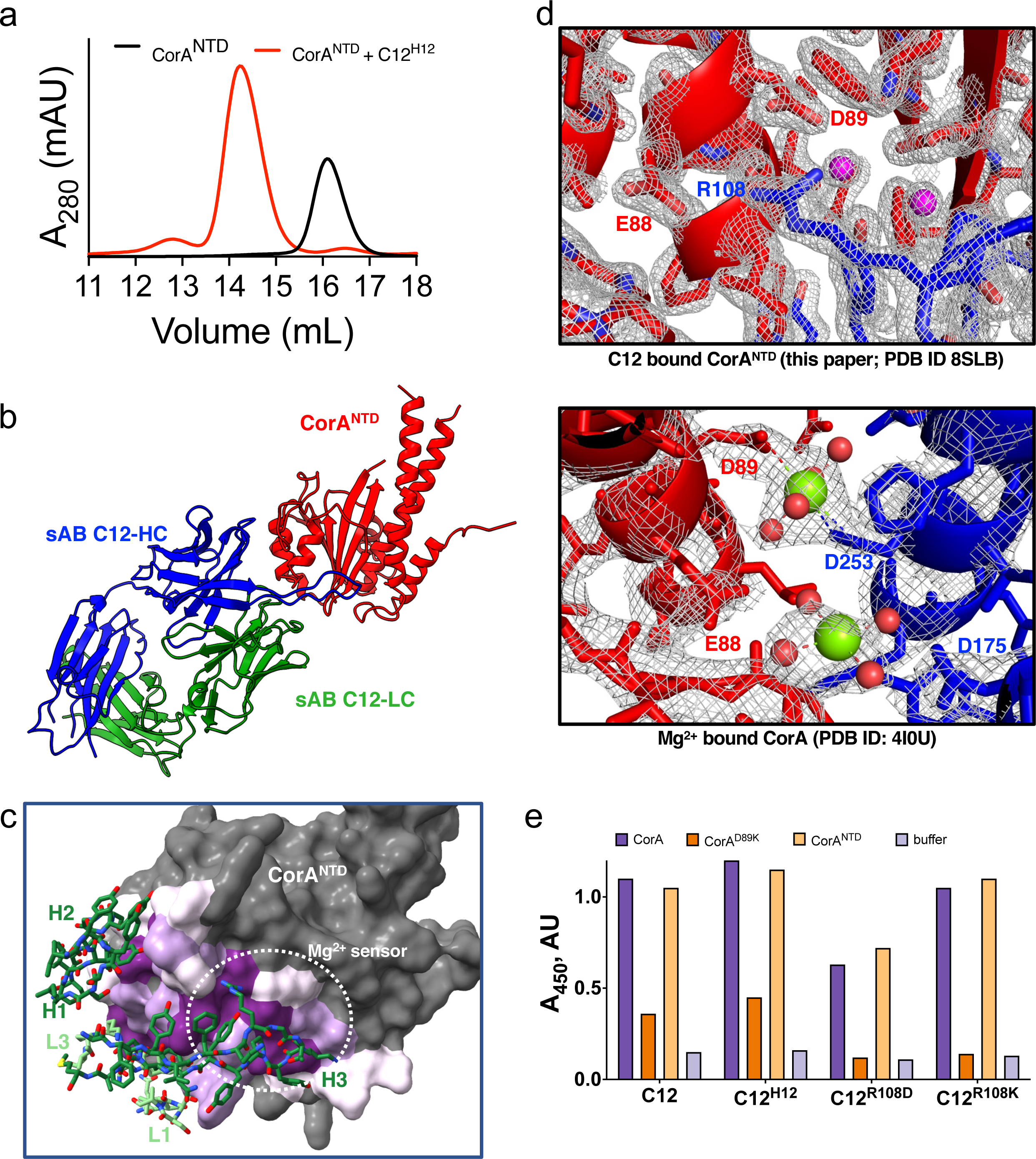
Structural analysis of the sAB C12 epitope. **A**. A stable complex of CorA^NTD^-C12^H12^ was isolated by SEC, with the complex chromatogram (red) showing a shift to an earlier retention volume compared with CorA^NTD^ alone (black). **B**. Crystal structure shows a 1:1 complex formed by CorA^NTD^ and sAB C12^H12^. CorA^NTD^ is red, sAB C12^H12^ heavy chain is blue, and sAB C12^H12^ light chain is green. See **Table 1** for data processing statistics. **C.** Interaction interface of sAB C12 with CorA^NTD^, colored by percentage buried surface area: 10-49%, light purple; 50-70%, medium purple; 71-100% dark purple. Non-interacting areas colored dark gray. C12 CDRs that are part of interface are colored by chain (heavy chain – dark green; light chain – light green). **D**. *Top panel* - A closer inspection of the epitope reveals a salt bridge interaction formed by sAB C12^H12^-R108 and CorA^NTD^-D89. D89 forms part of the magnesium sensor site. *Bottom panel* – In the magnesium-saturated closed crystal structure (PDB ID: 4I0U), D89 and D253 from adjacent monomers (shown as blue and red) form a bidentate interaction to bind a hydrated Mg^2+^. 2F_c_-F_0_ electron density maps, shown as light gray mesh, were contoured at 2.0σ and 1.5σ, respectively, for the CorA^NTD^-C12 and 4I0U structures. **E**. Binding analysis of sAB C12 to CorA reveals that the binding can be abolished by specific mutation of either CorA^NTD^ D89 or sAB C12 R108, indicating that this interaction contributes to magnesium-dependency of sAB binding to CorA.

**Table 1.**
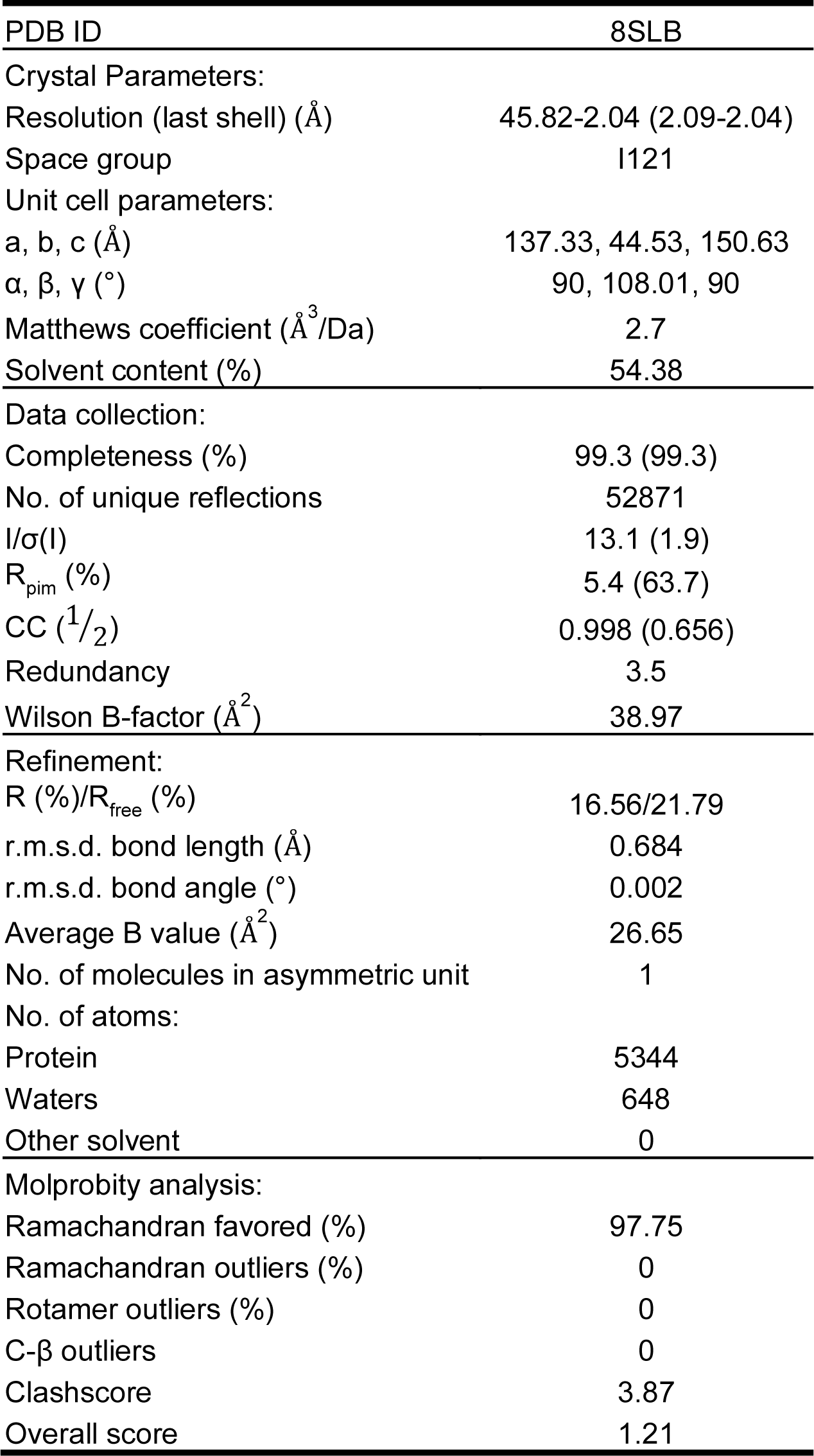
X-ray data collection, refinement, and validation statistics from Phenix[43]; r.m.s.d, root-mean-square deviation.

In order to crystallize this complex, we utilized an engineered variant of the sAB hinge region, known as H12, that had previously been shown to increase the crystallizability of Fab fragments based on the Herceptin scaffold[38]. C12^H12^ bound with identical affinity to CorA (**Supplemental Fig. 2C**). High quality crystals were obtained and we determined the structure of the CorA^NTD^-C12^H12^ complex at 2.0 Å resolution (**Fig. 4B, Table 1**). The crystal structure showed a one-to-one interaction between sAB C12 and CorA^NTD^. The overall architecture of the CorA^NTD^ complex with sAB C12 is highly similar to the structure of isolated CorA^NTD^ reported previously[28]. There is clear electron density for the core of the NTD domain where the divalent cation sensor sites are. The portions of CorA^NTD^ distal to these sites, which include portions of the α5, α6, and α7 helices, are frayed compared to their conformations in pentameric closed structure. This fraying results in density missing for the loop, spanning residues 197-206, connecting the α5 and α6 helices. The poor density in this region is similar to previous isolated N-terminal domain structures, and most likely caused by the absence of interactions present in the full-length protein[28]. Similarly residues 115-121, which form a β-hairpin that flanks the aforementioned cytoplasmic helices, are missing in our structure.

All complementarity-determining regions (CDRs) of sAB C12^H12^ are well-resolved. The interaction interface between sAB C12 and CorA^NTD^ encompasses about 1,200 Å^2^ with 350 Å^2^ contributed by the light chain (LC), and 850 Å^2^ by the heavy chain (HC) of sAB C12 (**Fig. 4C, Supplemental Table 2**). The light chain interacts primarily through CDRs L1 and L3, with L2 forming direct contacts only with the HC CDR-H3. The heavy chain interacts through all three CDRs, with the major contacts formed by CDR-H3. The structure revealed C12 has an extremely long CDR-H3, a 19-residue loop that forms an elongated anti-parallel β-sheet and interacts extensively with magnesium-binding residues of the cytosolic domain. Despite the crystals being grown in the presence of 20 mM MgCl_2_, neither the M1 nor M2 regulatory sites were occupied by the divalent cation. In the full-length CorA crystal and cryo-EM structures solved in the presence of this MgCl_2_ concentration, the sensor sites are all bound with magnesium ions[26–31,33].

A closer inspection of the interaction interface reveals ionic interactions between sAB C12 CDR-H3 residue R108 and CorA^NTD^ residues D89 and E88 (**Fig. 4D – top panel**). CorA residue D89 comprises one half of the aforementioned M1 cytosolic sensor, with the other half contributed by D253 of the adjacent protomer in the full-length CorA. This residue pair, along with the triad of residues, which includes E88, that principally forms the M2 site, is responsible for binding Mg^2+^ ions crucial to the cation-dependent gating and closing of the channel (**Fig. 4D – bottom panel**). The complex structure shows that the guanidinium group of R108 in sAB C12 effectively replaces Mg^2+^ in its binding pocket through formation of a salt bridge with D89 of CorA^NTD^. Indeed, when D89 is mutated to lysine (CorA^D89K^), the binding of C12 is significantly reduced, establishing a pivotal role for this interaction in the complex between CorA and the sAB (**Fig. 4E**). Similarly, when R108 is mutated to aspartate (C12^R108D^), the CorA-C12 interaction is diminished, but not when R108 is mutated to lysine (C12^R108K^). These results highlight the importance of both this ionic interaction and the integrity of the sensor as key determinants of sAB specificity. We conclude sAB C12 is conformation-specific by acting as competitive inhibitor of magnesium binding to the divalent cation sensor.

We superimposed the structure of CorA^NTD^-C12^H12^ to model sAB C12 binding to the full-length channel using both the closed (PDB ID: 4I0U) and open (PDB ID 3JCG) structures. Notably, the sAB epitope is a major contact point in the closed state of the channel between adjacent monomers of the pentameric assembly. When this section of CorA^NTD^ is superimposed onto the position of one of the protomers in the closed structure of CorA, significant steric impediments to sAB C12 binding in the context of the full-length pentameric protein are apparent (**Fig. 5A**). This suggests that C12 binds to conformations of full-length CorA that are quite distinct from the closed symmetric structures determined by X-ray crystallography and cryo-EM analyses done in high Mg^2+^ concentrations[26–31,33]. Indeed, when we superimposed the complex to the open state, these steric impediments are no longer present (**Fig. 5B**). These models make evident that the concerted movements of adjacent protomers to adopt the open state conformations are necessary to accommodate binding of C12. C12 CDR-H3 then forms a wedge between the protomers, and the models visually allude to sAB interactions potentially spanning the interfacial region between CorA subunits. Relatedly, we observed significant increases in both the on- and off-rates of C12 to CorA^NTD^ by SPR, with the latter supporting potential additional interactions with the full-length CorA contribute to the overall higher binding affinity (**Supplemental Fig 2D**, **Supplemental Table 1**). While we cannot model these interactions based on this preliminary analysis, it is possible a sAB raised against the full-length protein would recognize an epitope spanning a surface exposed only in the Mg-dep conformations. Indeed, binding data of sAB C18, which recognizes full-length CorA in the Mg-dep state but not the isolated N-terminal domain, provide evidence of precisely this scenario. Further, given C12 binds CorA across a broad range of magnesium concentrations, it is likely the sAB recognizes conformations distinct from both the open, Mg-dep states and the closed, symmetric, Mg^2+^-saturated states.

**Figure 5.**
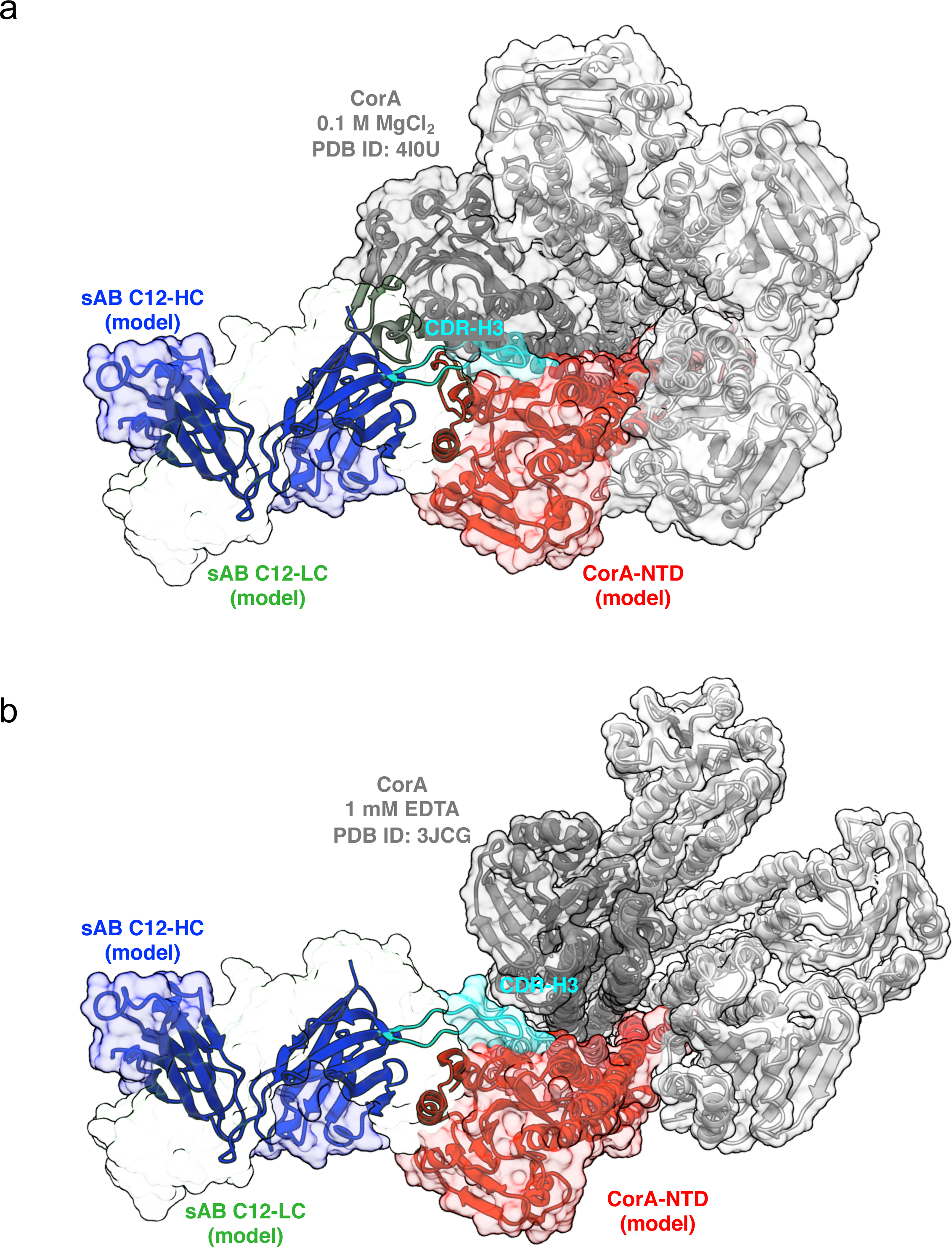
–. Modeling of the sAB C12 paratope-epitope interaction against the full-length CorA structure. In both figures the sAB light chain is shown as as transparent surface for clarity. A. Superposition of CorANTD (red) and sAB C12 (heavy chain – blue) onto the closed state pentamer (PDB: 4I0U; epitope-adjacent monomer colored dark gray, others light gray) reveals significant steric hindrance of C12 CDR-H3 (cyan) which precludes its binding to closed state of the channel. B. Superposition of CorANTD and sAB C12 (heavy chain colored as in A) onto the open state pentamer (PDB: 3JCG; monomers colored as in A) reveals complete accessibility of the epitope in the Mg-dep condition, which the sAB was selected in.

### CorA-C12 complexes adopt conformations similar to the magnesium depleted state

To evaluate the conformation of CorA in the presence of sAB C12, we utilized site-directed spin-labeling (SDSL) and electron paramagnetic resonance (EPR) spectroscopy. Single-cysteine mutants were introduced to the cysteine-free background[34]. Five different positions, all located within the cytoplasmic N-terminal domain, were chosen to report on the conformational changes of the channel between the Mg-dep and Mg-phys conditions: E191C, V199C, E206C, K215C, and S233C (**Fig. 6A**). E191 and V199 are located on the cytoplasmic α5-helix, and E206 is at the end of the negatively-charged glutamate-enriched loop connecting the α5-and α6-helices. K215 and S233 are both located on the α6 helix. These helices form the outer wall of the intracellular portion of the channel, while the glutamate-enriched loop is proximal to the basic sphincter formed by lysine residues at the C-termini of transmembrane helix 2 (TM2)[28]. Additionally, the M2 sensor site residues D175 and D179 are located on the α5-helix; due to this combination of features, this region was expected to exhibit sensitivity to Mg^2+^-dependent conformational changes (**Fig. 6A – right panel**) [29,33].

**Figure 6.**
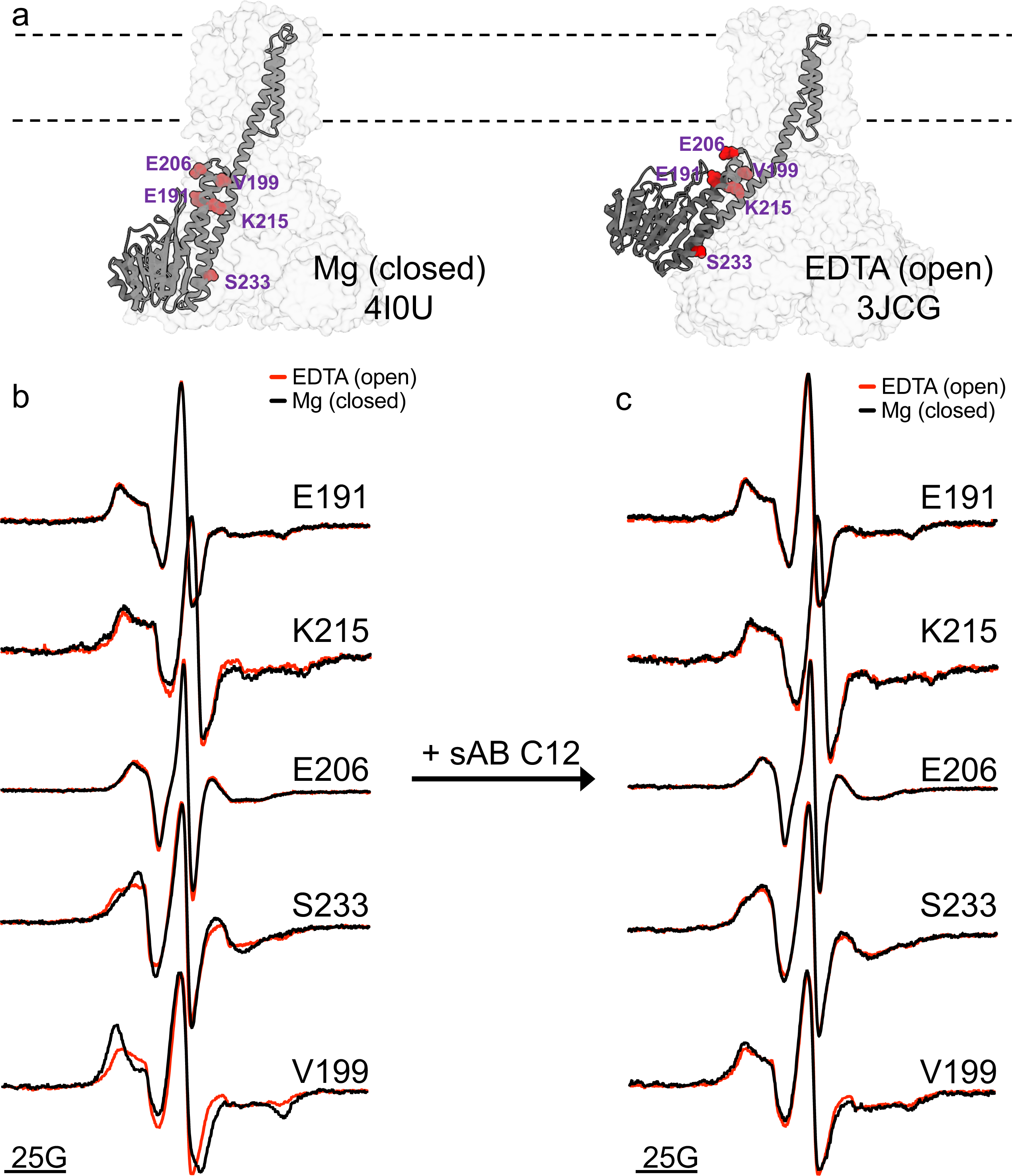
–. EPR spectroscopy analysis of CorA and CorA-C12 complex. **A.** Visualization of positions mutated to prepare single-cysteine CorA mutants for site-directed spin-labeling EPR spectroscopy. CorA structure 4I0U was used to visualized Mg^2+^-bound closed state (left). CorA structure 3JCG was used to visualized Mg^2+^-free form (right). Sidechains were added to 3JCG using Coot[42] purely for visualization purposes. Residues at indicated positions represented as red spheres and labeled accordingly. Mutated positions shown only for chain A, which undergoes significant conformational changes in the Mg^2+^-free “open” condition designated “State 1”, for clarity and illustrative purposes. However, all five possible positions for each mutant are labeled. Dotted lines denote boundary of transmembrane region. **B**. EPR spectra for all five single cysteine mutants of CorA in absence (1 mM EDTA, red) and presence of Mg^2+^ (black). Spectra are overlaid to show differences between the two conditions. The labeled mutant at V199C (bottom-most spectra) showed the most sensitivity to the two different conditions. **C.** EPR spectra for all five single cysteine mutants of CorA in absence (1 mM EDTA, red) and presence of Mg^2+^ (black) after the addition of sAB C12. Spectra are overlaid to show differences. The V199C mutant produced identical spectra (bottom-most spectra) in the two conditions after addition of sAB C12.

Each single cysteine mutant was conjugated with the thiol-specific methanethiosulfonate (MTSL) spin label. Continuous wave EPR (cw-EPR) data were collected for CorA and CorA-C12 in the presence and absence of Mg^2+^. For CorA alone, spin labeled CorA mutants E191C and K215C gave rise to similar spectra in the absence (1 mM EDTA) and presence of Mg^2+^ **(Fig. 6B – first two spectra)**. While the positions E206C and S233C showed subtle but significant spectral changes between the two conditions, the EPR data established the V199C position as the most sensitive to changes in Mg^2+^ concentrations (**Fig. 6B – last three spectra**). EPR spectra exhibited line-shape changes characteristic of increased mobility of the MTSL in EDTA relative to Mg^2+^ for all three of these mutants, suggesting a less restricted local environment. These changes are particularly evident in the low-field spectral lines. We infer from these data together with the low-resolution cryo-EM structures of the open state that the spectra report on the outward rotation of CorA protomers due to the depletion of Mg^2+^ from the regulatory sites (**Fig. 6A-B**)[33]. These data also established the utility of these mutations as positions capable of reporting on magnesium ion-induced conformational changes in CorA in EPR spectroscopic studies of the channel.

We then recorded spectra of the CorA-C12 complex either in the presence or absence of Mg^2+^. While 191 and 215, as expected, gave rise to no differences in either set, EPR spectra for 199, 206, and 233 all showed virtually identical line-shapes between the EDTA and Mg^2+^ conditions (**Fig. 6C**). The spectral lines in both conditions showed increased mobility of the MTSL characteristic of the EDTA state. This was most pronounced in CorA^V199C^, and thus consistent with the CorA alone EPR data (**Fig. 6C, bottom-most spectra**). These data suggest sAB C12 prevents the Mg^2+^-dependent conformational changes typically observed at the divalent cation sensors in the presence of 20 mM MgCl_2_, resulting in the closed, magnesium bound state. These observations are consistent with our complex formation and competition ELISA data (**Fig. 3**), which demonstrate much higher concentrations of Mg^2+^ are required to completely displace sAB C12 from CorA to produce the closed, Mg^2+^-saturated form of the channel. Combined with the structural data and the mapping of the sAB epitope onto the closed symmetric and open asymmetric states of the pentameric channel (**Fig. 4-5**), C12 apparently stabilizes an open-like conformational state even in the presence of 20 mM Mg^2+^ by competing with divalent cations for binding to the cytoplasmic regulatory sites.

## Discussion

CorA is a prototypical ion channel that is regulated through a mechanism that involves a series of ion dependent conformational transitions[34–36]. Previous studies of this channel had suggested that at high concentrations of Mg^2+^ the channel exists as a symmetric pentameric structure with a constricted pore region that prevents ion permeability[29]. In the absence of Mg^2+^, the channel becomes highly asymmetric, adopting multiple conformational states. However, because of the conformational heterogeneity, the resolutions of these structures were low and these conformations could not be characterized with confidence[32,33]. There are thus many outstanding questions pertaining to how CorA is regulated and the structural roadmap for how Mg^2+^ is ultimately transported through the channel.

In order to gain insight into these fundamental questions, we exploited a phage display selection strategy developed by us that allows the generation of conformational specific sABs[2,4,7]. The biopanning conditions are designed to stabilize specific functional states of proteins which operate through programmed conformational transitions. We had shown in previous work that this approach has been highly successful in stabilizing functionally relevant forms of proteins that are too transient to be studied by traditional methods[3,8,9]. Notably, it is important to emphasize that sABs that are generated to stabilize particular conformations by our selection strategy are opportunistic, not deterministic of that conformation. That is, they are selected through a binding mechanism akin to conformational selection, not induced fit.

From a cohort of sABs that were generated from CorA selections done in the absence of Mg^2+^, we selected for further study two sABs (C12 and C18) that displayed different degrees of Mg^2+^ sensitivity. sAB C12 binds CorA in both Mg^2+^ depleted and physiological Mg^2+^ conditions, while C18 is highly selective to the depleted condition. Size exclusion chromatography and negative staining analysis in Mg^2+^ depleted conditions showed that the CorA complexes in the case of both sABs existed in either 1:1 or 2:1 sAB-CorA stoichiometries, albeit at different ratios. C12-CorA resides in a 1:1 stoichiometry at high [Mg^2+^], and decreasing the concentration converts it to a largely 2:1 complex. C18-CorA exists in 2:1 and 1:1 forms in the absence of Mg^2+^; this reflects the identification of at least two different conformations in the CorA cryo-EM structures in 1 mM EDTA[33]. This suggests that in these open-like conditions there are two accessible epitopes for C18 binding in context of the full pentameric channel; however, the epitope to which C12 binds has different accessibility that supports a stepwise binding process. Thus, these two sABs probe different features of the asymmetric channel under open-like conditions. Notably, it is clear that the conformational forms in this condition do not support binding of more than two copies of the same sAB simultaneously.

A by-product of generating conformationally specific sABs is that they can be used as a gauge to assess the energy landscape involved in transitions from one functional state to another[23]. For instance, for the conformationally specific sABs that were selected in the Mg^+2^ depleted conditions, determining the change of their binding affinities as a function of [Mg^+2^] provides a direct measure of the ΔΔG coupling between the two states with respect to their particular epitopes. For the highly [Mg^+2^] dependent C18 sAB, the binding affinity differences between 1 mM EDTA and 1 mM MgCl_2_ conditions is over 3 orders of magnitude (4.1 kcal/mol). In contrast, even though the sABs were obtained from the same selection pool, sAB C12 exhibits far less [Mg^+2^] sensitivity, indicating it sits on an epitope that is less affected by the underlying energetics controlling the ion dependent conformational transitions. However, this does not mean that the C12 epitope is outside the influence of the conformational changes occurring during ion binding. On the contrary, the C12-CorA^NTD^ structure indicates the sAB should bind between two CorA oligomers, effectively capturing a highly asymmetric structure that resembles an open-like conformation based on the EPR spectroscopic data. Given the sAB was raised against CorA in 1 mM EDTA, we interpret the structural, biophysical, and biochemical data to mean that the full-length pentameric channel presents an epitope to C12 that is essentially equally accessible at both depleted and physiological [Mg^+2^] levels. In contrast, the C18 epitope becomes less available at increasing [Mg^+2^] as suggested by the markedly decreased on-rates observed from the kinetic data. A comprehensive structural accounting of the location of these two different epitopes awaits the high resolution analysis of these sABs bound to the pentameric channel, which is currently ongoing.

## Materials and Methods

### Protein purification and nanodisc reconstitution

CorA, CorA^NTD^, all mutants of both, and the sABs were expressed and purified as previously described[7,14,38], with full-length CorA and CorA^NTD^ purified in the presence of 20 mM MgCl_2_ to prevent binding of other cations during purification. Mutations for CorA and the sABs were introduced by site-directed mutagenesis and sequence verified at the University of Chicago Comprehensive Cancer Center Sequencing Facility. Lipids for nanodiscs were prepared by combining chloroform solutions of POPC and POPG (Avanti) in a 4:1 molar ratio, evaporating the chloroform using a stream of nitrogen gas to produce a thin lipid film, and drying overnight under vacuum. The dried film was then resuspended in buffered water with 2% (w/v) β-n-dodecylmaltoside (Anatrace) and sonicated (Branson sonifier) to produce a clear solution of lipid:detergent micelles. Nanodisc reconstitution was adapted from a previously described protocol for CorA[7,33], using MSPΔH5 instead of MSP1D1, and a CorA:MSP:lipid molar ratio of 1:10:350, where one mole of CorA is defined as the ion channel pentamer. Nanodiscs were formed by adding 800 mg of Bio-beads SM2 (Bio-rad) per 1 mL of reconstitution mixture and incubating overnight at 4°C with nutation. CorA-loaded nanodiscs were separated from empty ones by size-exclusion chromatography (SEC) using a Superdex200 Increase 10/300 column (Cytiva). For complexes with sAB C12, CorA nanodiscs were purified in Hepes-buffered saline (HBS) with different concentrations of MgCl_2_, while for complexes with sAB C18, CorA nanodiscs were purified in HBS with 1 mM EDTA. The complex of CorA^NTD^ and sAB C12^H12^ was similarly purified by SEC in HBS containing 20 mM MgCl_2_. For long-term storage, nanodiscs were supplemented with 10% (v/v) sucrose. All proteins and nanodisc preparations were stored in aliquots at –80°C and thawed freshly prior to each experiment, unless stated otherwise.

### Characterization of sAB binding by single-point and competition ELISA

Single-point ELISA was performed to measure the binding of purified wild-type and mutant sABs using constant concentrations of 200 nM. Competition ELISA were performed to estimate CorA affinity for Mg^2+^ in the presence of constant concentration of sAB C12. For both sets of experiments, the CorA sample was prepared as follows. CorA was reconstituted into nanodiscs as above, except using chemically biotinylated MSPΔH5, which was biotinylated using the NHS-PEG4-biotin reagent (Thermo) as previously described[4], and CorA-loaded nanodiscs were purified by size-exclusion chromatography using HBS with 1 mM EDTA. CorA nanodiscs were then diluted into ELISA buffer (HBS with 2% BSA) to a final concentration of 25 nM. Prior to ELISA, high-binding 96-well plates (Nunc) were coated with 2 µg/mL neutravidin and blocked with buffer containing 1% BSA. Plates were washed three times with ELISA buffer, and CorA was allowed to bind to wells for 30 minutes. Following immobilization, wells were washed three times with ELISA buffer.

For single-point ELISAs, 200 nM of each sAB was diluted into ELISA buffer containing either 1 mM EDTA or 20 mM MgCl_2_. The sAB solutions were then added to wells with either CorA immobilized or buffer alone for background measurements. Following a 30-minute incubation, wells were washed three times with ELISA buffer. An HRP-conjugated goat anti-human secondary antibody (Jackson Immunoresearch) was then added and incubated for 30 minutes. Wells were washed again three times with the appropriate buffers and bound sABs were detected with TMB substrate (Thermo).

For competition ELISAs, 50 nM of each sAB was diluted into ELISA buffer containing increasing concentrations of MgCl_2_, from 0 to 250 mM. The sAB/MgCl_2_ solutions were then added to wells with either CorA immobilized or buffer alone for background measurements. Following a 30-minute incubation, wells were washed three times with ELISA buffer containing the appropriate MgCl_2_ concentration for each experiment. An HRP-conjugated goat anti-human secondary antibody (Jackson Immunoresearch) was then added and incubated for 30 minutes. Wells were washed again three times with the appropriate buffers and bound sABs were detected with TMB substrate (Thermo).

All experiments were performed at room temperature and done in triplicate. Data were analyzed in GraphPad Prism 7 and, where applicable, EC_50_ values were calculated using a variable slope model with the assumption of sigmoidal dose response.

### Binding kinetics by surface plasmon resonance

The interaction between sABs C12 and C18 and CorA was further dissected by SPR. Analysis of C12 binding to CorA was performed using a BIACORE 3000 (Cytiva) by immobilizing his-tagged DDM-solubilized CorA onto a nitrolotriacetic acid sensor chip. A 2-fold dilution series of C12 (25 nM to 1.6 nM) was injected at a flow rate of 30 μL/min, using a 180 seconds association phase and 600 seconds dissociation phase. C12 binding was measured both in the absence (SPR buffer: 10 mM Hepes, 150 mM NaCl, 50 µM EDTA, 0.05% DDM) and in the presence of Mg^2+^ (SPR buffer with 20 mM MgCl_2_). All sensorgrams were double-reference subtracted using a channel without CorA immobilized and with buffer-only injections.

Analysis of C18 binding to CorA was performed using a MASS-1 instrument (Bruker) with a His-capture sensor chip (XanTec). Experimental and reference channels were prepared as described above. A 2-fold dilution series of C18 (200 nM to 12.5 nM) was prepared in SPR buffer supplemented with MgCl_2_ when required, and injected as above, using a 180 seconds association phase and 300 seconds dissociation phase. C18 binding was measured in the following MgCl_2_ concentrations: 0, 250 µM, 500 µM, 750 µM, 1 mM, 1.5 mM, and 2 mM. Sensorgrams were double-referenced as above and, except for the experiments in 1.5 and 2 mM MgCl_2_ that showed no binding, analyzed as described below.

For all SPR experiments, data processing and kinetic analysis were performed using the BiaEvaluation software suite (Cytiva). All data sets were fit to a 1:1 interaction model to determine kinetic rate constants. Double-reference subtracted sensorgrams and the kinetic fitted curves were exported and plotted using Prism 7.0 (GraphPad). Differences in the change of Gibbs free energy (ΔΔG^0^) was calculated from the dissociation constants (K_D_) at two different MgCl_2_ concentrations[8].

### Negative stain EM of CorA-C12 and CorA-C18 complexes

To determine sample quality and stoichiometry of sAB binding, complexes of CorA-C12 in either 1 mM EDTA, 12.5 mM MgCl_2_, or 40 mM MgCl_2_, and CorA-C18 in 1 mM EDTA were isolated by SEC as described above. Purified proteins were diluted to 0.005-0.01 mg/mL and 3.5 µL was applied onto copper grids with a thin layer of continuous carbon (approximately 3-4 nm, Electron Microscopy Sciences) that had been plasma-cleaned (Gatan Solarus) using air for 30 seconds. Excess protein solution was blotted (Whatman 4) and grids were washed once using a water droplet, after which 3.5 µL of freshly-prepared 1% uranyl formate was applied, immediately blotted, and added again. After 30 seconds, uranyl formate was blotted and grids were dried in air. Grids were imaged using a Tecnai F30 microscope (FEI) with a 4k x 4k digital CCD camera at a pixel size of 2.3 Å per pixel. In total, 50-100 images were collected per dataset. All images were processed using Relion software (version 3.1) to obtain two-dimensional class averages[39]. Image analysis was additionally performed using the ImageJ software.

### Crystallization, data collection, and refinement

Initial efforts to obtain the structure of the N-terminal domain of CorA in complex with wild-type sAB C12 were unsuccessful, as this complex did not yield crystals that diffracted past 10 Å. A mutation shown to improve the crystallizability of the 4D5 scaffold, FNQIKG, was introduced into sAB C12, resulting in sAB C12^H12^,[38]. Purified CorA^NTD^ was mixed with a 1.2-fold molar excess of sAB C12^H12^ and the complex was isolated by size exclusion chromatography (Superdex 200 10/300 GL, Cytiva) in 25 mM HEPES (pH 7.5), 200 mM NaCl, and 20 mM MgCl_2_. Fractions containing the purified complex were pooled, concentrated, and immediately used for crystallization.

Crystallization trials of the CorA^NTD^-C12^H12^ complex were set up using a Mosquito robot (SPT Labtech) and the PEG/Ion Screen (Hampton Research). Protein solutions containing 18.7 mg/mL of the complex were mixed with equal volumes of reservoir solution and grown at room temperature by the hanging drop vapor diffusion method. Initial clusters of needle-like crystals were obtained from condition H5: 16% PEG 3350 and 0.02 M citric acid/ 0.08 M bis-Tris propane pH 8.8. These were then optimized using the Additive Screen (Hampton Research), with condition E3 (3% dextran sulfate in 16% PEG 3350 and 0.02 M citric acid/ 0.08 M bis-Tris propane pH 8.8) resulting in well defined, larger needle-like crystals. Finally, a last round of optimization was performed around this condition, resulting in rod-like crystals grown in 8% dextrane sulfate, 17% PEG 3350, 0.02 M citric acid/0.08 M bis-Tris propane pH 8.8. These crystals were harvested, cryoprotected in 8% dextrane sulfate, 17% PEG 3350, 0.02 M citric acid/0.08 M bis-Tris propane pH 8.8, 20% ethylene glycol, and subsequently flash frozen in liquid nitrogen.

A dataset was collected from a single crystal at 100 K at beamline 23ID-B at the Advanced Photon Source. The dataset was processed, scaled, and merged using XDS[40]. Data collection and processing statistics are listed in **Table 1**. The CorA^NTD^-sAB C12^H12^ complex structure was determined to 2.0 Å by molecular replacement with PHASER, using the previously determined crystal structures of a sAB (PDB: 3PGF) without the complementarity determining regions (CDRs) and CorA^NTD^ (PDB: 2BBH) as the search models[41]. Iterative cycles of manual model building and refinement with COOT, PHENIX, and REFMAC were used to build the final complex model[42–44]. Progress of refinement was monitored by steady decreases in the R and R_free_ values. Model geometry validation was performed using Molprobity[45].

Protein surface interactions were calculated using the Proteins, Interfaces, Structures, and Assemblies server[46] and visualized using PyMol (Schrödinger, LLC). All other images were prepared using Chimera and ChimeraX[47,48]. The crystal structure was deposited into the Protein Data Bank (PDB ID 8SLB).

### Continuous-wave electron paramagnetic resonance (EPR) spectroscopy

CorA cysteine mutants were expressed as previously described[34]. After elution from the immobilized metal affinity chromatography (IMAC) column, purified CorA was reacted with a 10-fold molar excess of 1-Oxyl-2,2,5,5-tetramethylpyrroline-3-methyl)methanethiosulfonate (MTSL; Toronto Research Chemicals) for 30 min on ice, followed by a second addition of 10-fold molar excess of MTSL and an additional 30 min reaction. The reaction was stopped by the addition of 1000-fold excess of L-Cysteine and the excess probe was removed by size exclusion chromatography. Reconstitution was carried out by destabilizing preformed POPC-POPG (3:1 w/w) liposomes with a saturating concentration of Triton™ X-100 (0.2 w/w ratio) before adding labeled CorA to the liposome suspension in a 1/1500 protein/lipid molar ratio. Detergent was removed by successive addition of Bio-Beads SM-2 (Bio-Rad) and proteoliposomes were harvested by ultracentrifugation at 200,000g and resuspended into 200 µL of buffer. The samples were divided into two portions and each was equilibrated with buffer containing either 1 mM EDTA or 20 mM MgCl_2_. Continuous-wave EPR (cw-EPR) spectroscopic measurements were performed at room temperature on a Bruker EMX X-band spectrometer equipped with a dielectric resonator (ER 4123D). Spin labeled samples were loaded into a gas-permeable TPX plastic capillary and spectra were recorded at 2.0 mW incident power and 100 kHz modulation frequency with 1.0 G of modulation amplitude. Prior to and during the recording of EPR spectra, the samples were equilibrated with N_2_.

## Acknowledgements

We thank Ekaterina Filippova for assistance with structure refinement and deposition, Somnath Mukherjee for help in setting up and interpreting SPR experiments, and Duy P. Hua for advice and critiques in the preparation of this manuscript. We express our thanks to Lucas J. Bailey for advice on the use of sAB elbow variants to enhance crystallizability. We thank the staff at the University of Chicago Advanced Electron Microscopy Facility and the University of Chicago Comprehensive Cancer Center Sequencing Facility for their support. We thank the staff at the Advanced Photon Source Beamline 23-ID at Argonne National Labs for their assistance in the collection of diffraction data. This project was funded by NIH grants R01GM117372 (to A.A.K.) and R01GM120561 (to E.P.).

## Author Contributions

S.K.E., P.K.D., O.D., E.P., and A.A.K. conceived the project. P.K.D., D.D, and O.D. generated the sABs. S.K.E., P.K.D., D.D., B.M.S., and P.T. prepared samples and performed sAB validation. S.K.E. and P.T. performed negative-stain EM experiments. P.K.D, D.D, S.S.K., B.G.R., and S.K.E. performed crystallization and structure determination. P.K.D., S.S.K., O.D., and E.P. performed EPR experiments. S.K.E., P.K.D., and A.A.K. wrote the manuscript with input from the other authors.

## Data Availability

Atomic coordinates for the crystal structure of CorA N-terminal domain in complex with sAB C12 have been deposited in the Protein Data Bank under accession number 8SLB.

---

**Supplemental Figure 1.**
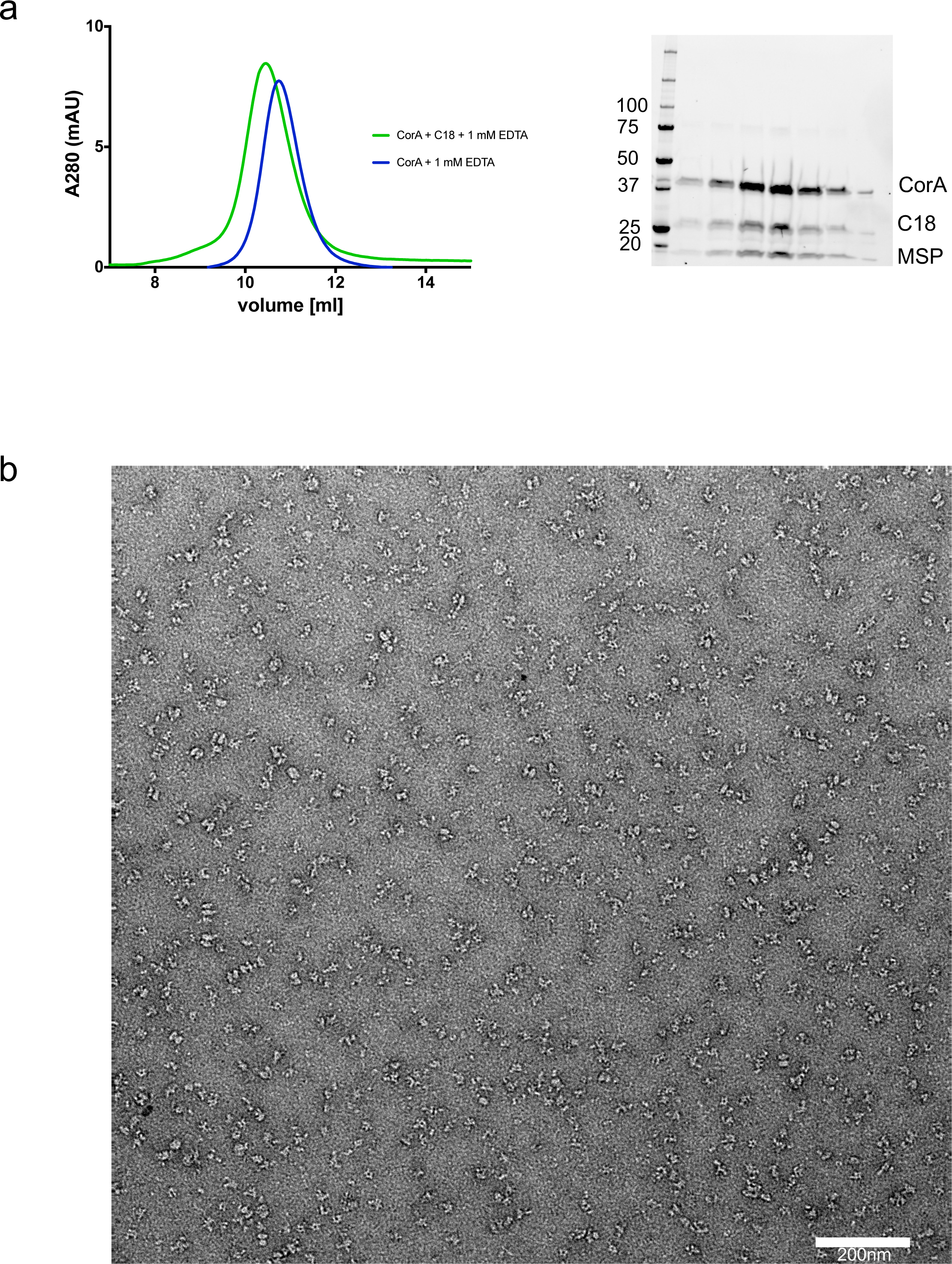
Analysis of complex formed by sAB C18 with nanodisc-reconstituted CorA in 1 mM EDTA. **A**. SEC chromatograms of CorA-C18 (green) and CorA alone (black) overlaid together. SDS-PAGE gel image of the complex is shown on the right. **B**. Negative-stain EM sample micrograph of CorA-C18 complex.

**Supplemental Figure 2.**
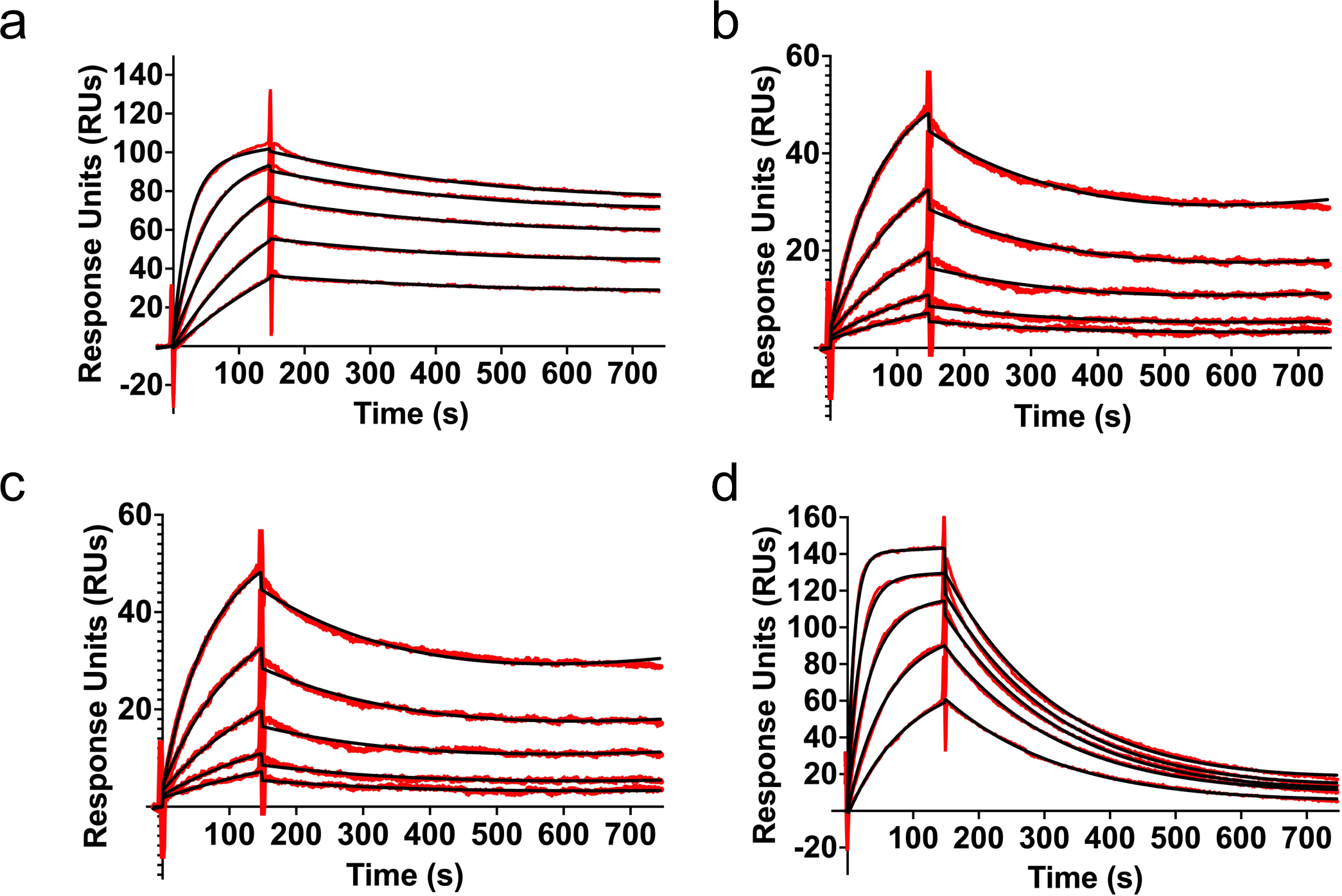
Interaction of C12 with CorA and CorA^NTD^. **A**. Double-referenced sensorgrams (red) of a 2-fold dilution series of C12 (25 – 1.56 nM) binding to CorA in the absence of Mg^2+^ overlaid with kinetic fitted curves (black) from a one-to-one binding model. **B**. The same as in A., except the experiment was performed with 20 mM MgCl_2_ in the running buffer. **C**. The same as in B, except sAB C12^H12^ was used instead of the wt. **D**. The same C12^H12^ dilution series was tested using CorA^NTD^ instead of full-length CorA.

**Supplemental Figure 3.**
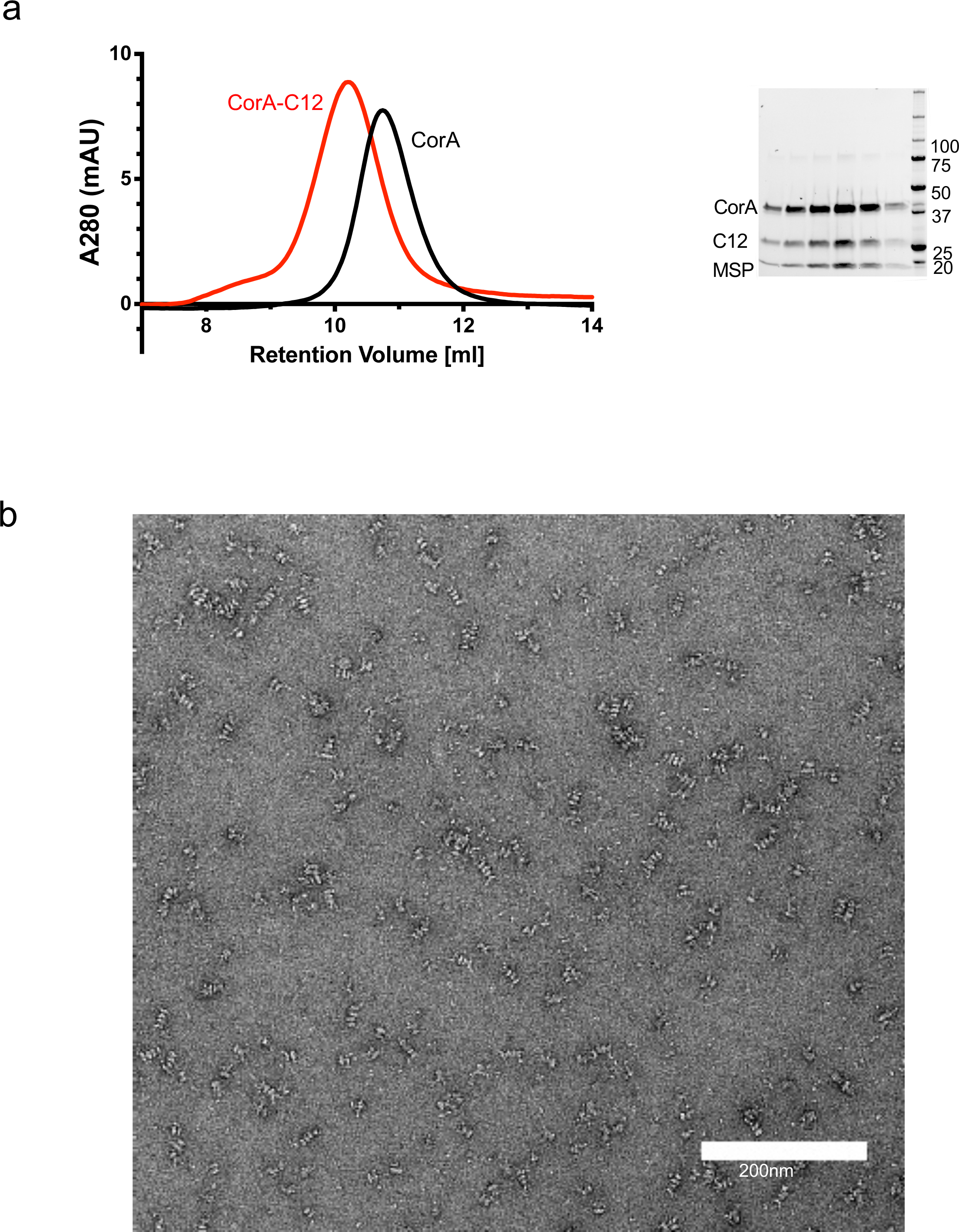
Analysis of complex formed by sAB C12 with nanodisc-reconstituted CorA. **A**. SEC chromatograms of CorA-C12 (red) and CorA alone (black) overlaid together. SDS-PAGE gel image of the complex is shown on the right. **B**. Negative-stain EM sample micrograph of CorA-C12 complex.

**Supplemental Table 1.** SPR kinetic parameters of sAB C12 and C18 binding to CorA in different environmental Mg^2+^ concentrations.

| Experiment | $k_{\text{on}}$ ($\text{M}^{-1}\text{s}^{-1}$) | $k_{\text{off}}$ ($\text{s}^{-1}$) | $K_{\text{D}}$ (nM) |
| --- | --- | --- | --- |
| C12 binding to CorA without $\text{Mg}^{2+}$ | $1.6 \times 10^6$ | $1.3 \times 10^{-3}$ | 0.83 |
| C12 binding to CorA with 20 mM $\text{Mg}^{2+}$ | $5.2 \times 10^5$ | $3.1 \times 10^{-3}$ | 5.9 |
| C12<sup>H12</sup> binding to CorA with 20 mM $\text{Mg}^{2+}$ | $5.1 \times 10^5$ | $3.1 \times 10^{-3}$ | 6.0 |
| C12 binding to CorA<sup>NTD</sup> without $\text{Mg}^{2+}$ | $3.6 \times 10^6$ | $5.3 \times 10^{-3}$ | 1.5 |
| C18 binding to CorA without $\text{Mg}^{2+}$ | $2.8 \times 10^5$ | $2.9 \times 10^{-3}$ | 11.0 |
| C18 binding to CorA with 250 $\mu\text{M}$ $\text{Mg}^{2+}$ | $1.9 \times 10^5$ | $5.2 \times 10^{-3}$ | 28.0 |
| C18 binding to CorA with 500 $\mu\text{M}$ $\text{Mg}^{2+}$ | $7.5 \times 10^4$ | $4.9 \times 10^{-3}$ | 65.8 |
| C18 binding to CorA with 750 $\mu\text{M}$ $\text{Mg}^{2+}$ | $5.2 \times 10^3$ | $6.5 \times 10^{-3}$ | 1,260 |
| C18 binding to CorA with 1.0 mM $\text{Mg}^{2+}$ | $1.5 \times 10^2$ | $1.7 \times 10^{-2}$ | 11,300 |
| C18 binding to CorA with 1.5 mM $\text{Mg}^{2+}$ | n.d. | n.d. | - |
| C18 binding to CorA with 2.0 mM $\text{Mg}^{2+}$ | n.d. | n.d. | - |

**Supplemental Table 2.** Hydrogen bonds and salt bridges between CorA^NTD^ and sAB C12^H12^.

| **CorA** | **C12** | **Distances (Å)** |
| --- | --- | --- |
| **N-terminal domain** | **Heavy Chain** |  |
| GLU88[OE1] | ARG108[NH1] | 2.8 |
| ASP89[OD1] | ARG108[NH1] | 3.3 |
| ASP89[OD2] | ARG108[NH1] | 4.0 |
| ASP89[OD1] | ARG108[NH2] | 3.1 |
| ASP89[OD2] | ARG108[NH2] | 2.9 |
| GLY15[O] | TYR55[OH] | 3.1 |
| GLN74[OE1] | SER58[OG] | 3.7 |
| LEU85[O] | ARG108[NH1] | 3.3 |
| GLY81[O] | TYR117[OH] | 2.7 |
| TYR105[OH] | TYR117[OH] | 3.7 |
| LEU17[N] | TYR55[OH] | 3.0 |
| PHE101[N] | LYS111[O] | 2.8 |
| HIS83[NE2] | ASN116[O] | 2.6 |
| HIS83[N] | TYR117[OH] | 3.5 |
|  | **Light Chain** |  |
| GLU103[OE1] | ARG67[NH1] | 3.0 |
| GLU103[OE1] | ARG67[NH2] | 3.6 |
| GLU103[OE2] | ARG67[NH1] | 3.5 |
| GLU103[OE2] | ARG67[NH2] | 2.5 |
| ASP55[OD1] | GLN28[NE2] | 3.6 |
| GLU78[O] | LYS95[N] | 3.0 |
| GLU103[OE1] | SER31[OG] | 2.7 |
| ASN104[OD1] | SER29[OG] | 3.6 |
| ARG54[NH2] | SER1[O] | 3.0 |
| ASN104[ND2] | SER29[OG] | 3.2 |

